# Identification of a *Nocardia seriolae* secreted protein targeting host cell mitochondria and inducing apoptosis in fathead minnow (FHM) cells

**DOI:** 10.1101/341651

**Authors:** Jianlin Chen, Wenji Wang, Liqun Xia, Zhiwen Wang, Yishan Lu, Jiahui Huang, Suying Hou

## Abstract

*Nocardia seriolae*, is a Gram-positive, partially acid-fast, aerobic, and filamentous bacterium. This bacterium is the main pathogen of fish nocardiosis. A bioinformatic analysis based on the genomic sequence of the *N*. *seriolae* strain ZJ0503 showed that *ORF3141* encoded a secreted protein with a signal peptide at the N-terminate which may target the mitochondria in the host cell. However, the functions of this protein and its homologs remain unknown. In this study, we experimentally tested the bioinformatic prediction on this protein. Mass spectrometry analysis of the extracellular products from *N. seriolae* showed that ORF3141 was a secreted protein. Subcellular localization of the ORF3141-GFP fusion protein revealed that the green fluorescence protein co-localized with the mitochondria, while ORF3141Δsig-GFP (with the signal peptide deleted) fusion protein was evenly distributed in the whole cell of fathead minnow (FHM) cells. Thus, the N-terminate signal peptide had a significant impact on mitochondrial targeting. Notably, the expression of ORF3141 protein changed the distribution of mitochondria from perinuclear halo into lumps in the transfected FHM cells. In addition, apoptotic features were found in the transfected FHM cells by overexpression of ORF3141 and ORF3141Δsig proteins, respectively. Quantitative assays of mitochondrial membrane potential value, caspase-3 activity and apoptosis-related gene mRNA expression suggested that cell apoptosis was induced in the transfected FHM cells. In conclusion, the ORF3141 was a secreted protein of *N*. *seriolae* that targeted host cell mitochondria and induced apoptosis in FHM cells. This protein may participate in the cell apoptosis regulation and plays an important role in the pathogenesis of *N*. *seriolae*.

**Author summary:** *Nocardia seriolae* is the causative pathogen responsible for fish nocardiosis. This facultative intercellular bacterium, adapts to survive and colonize by evading intracellular killing after being engulfed with macrophages in the host. Despite considerable economic losses caused by *N*. *seriolae* in fish infection, the pathogenic mechanism and specific virulence factor of this bacterium remain ambiguous. In this study, the characteristic of ORF3141 protein function was investigated by subcellular localization and its possible contributions on the ability of *N*. *seriolae* to induce apoptosis in transfected fathead minnow (FHM) cells was investigated. Here, we confirmed that ORF3141 was a secreted protein that targeted host cell mitochondria and induced cell apoptosis in FHM cells. Interestingly, after deleting the signal peptide, ORF3141Δsig protein was evenly distributed in the whole host cell and did not co-localize with the mitochondria which could also induce cell apoptosis. Thus, the N-terminate signal peptide played an important role in mitochondrial targeting, and the domain part without the signal peptide had a critical relationship with cell apoptosis. These results demonstrated that ORF3141 mays act as a potential virulence factor that induces apoptosis in fish cells. This protein is significant to elucidate the pathogenic mechanism of *N*. *seriolae* and this study mays provide beneficial insight to prevent and treat fish nocardiosis.

## Introduction

Fish nocardiosis is a systemic bacterial disease with three kinds of pathogenic bacteria, namely, *Nocardia seriolae*, *N. samonicida* and *N. asteroids*, which have been isolated from diseased fish [1–3]. The outbreak of fish nocardiosis is frequently reported in global aquaculture industries and has caused substantial commercial losses in Southeast Asia, especially China [4–7]. However, the pathogen of fish nocardiosis that has been reported with considerable frequency is *N*. *seriolae*. Notably, *N*. *seriolae* is the most frequently isolated *Nocardia sp*. from diseased fish [7]. This bacterium is an opportunistic pathogen, by infecting immunocompromised fishes through feeds, gills, and wounds and causing chronic systemic granulomatous disease [8]. The clinical signs of infected fish include skin ulcers and numerous white nodular structures on the gills and in the head kidney, trunk kidney, spleen, and liver [9]. *N*. *seriolae* infections have been documented in more than 39 kinds of fish, both freshwater and especially marine species, such as golden pompano (*Trachinotus ovatus*), snubnose pompano (*T. blochii*), yellow croaker (*Larimichthys crocea*), northern snakehead (*Channa argus*), blotched snakehead (*C*. *maculata*), largemouth bass (*Micropterus salmoides*), yellow tail (*Seriolae quinqueridiata*), amberjack (*S. dumerelli*) and sea bass (*Lateolabrax japonicus*) [5,10,11].

The pathogenic mechanisms and specific virulence factor of *N*. *seriolae* remain unknown. Beaman found that the virulence of *N. asteroides* was associated with its resistance to oxidative killing by phagocytes, inhibition of phagosome-lysosome fusion, neutralization of phagosomal acidification, and alteration of lysosomal enzymes within phagocytes, as well as the secretion of toxic substances by the nocardia, which are apparently involved in these processes [12]. Previous studies showed that *N. asteroides* might induce apoptosis in host cells [13–15]. The secretome studies of pathogenic actinomycetes have shown that secreted proteins, especially several mitochondrial targeted proteins, are closely related to their pathogenicity and may play an important role in the regulation of cell apoptosis and bacterial pathogenesis [16–19]. The modulation of cell apoptosis in the host by pathogenic bacteria is a major pathogenicity mechanism. Many bacterial proteins have a highly specific activity with the host mitochondria, which participate in the cell apoptosis by influencing the signaling pathways [20–22]. The mechanism of mitochondrial pathway regulation of apoptosis is very complex. The mitochondrial targeted protein allows the mitochondrial outer membrane permeabilization (MOMP) accompanied with the loss of mitochondrial membrane potential (ΔΨm). Subsequently, various apoptotic mediators (mammalian serine protease (Omi/HtrA2), second mitochondrial activator of caspase (SMAC), direct IAP binding protein with low pI (Diablo), cytochrome C (Cyt C), and apoptosis-inducing factors (AIF)) are released into the cytosol. The releases of Cyt c in cytosol causes the association of apoptosis protease-activating factor-1 (Apaf-1) and ATP/dATP to form apoptotic body, while AIF enters the nucleus acting in a caspase independent manner by causing chromatin condensation and fragmentation. Finally, the downstream procaspase-3 is activated, and cell apoptosis is induced in the host [23–25]. Thus, we speculated whether the mitochondrial targeting secreted proteins from *N*. *seriolae* are potential virulence factors.

A bioinformatic analysis based on the whole genome sequence data of *N*. *seriolae* strain ZJ0503 [26] showed that *ORF3141* encoded a secreted protein which might target the mitochondria in the host cell, and the ORF3141 protein had a signal peptide at the N-terminate (residues 1-30, MKLLNPRGFGLVCASAAVAAGLMLAGCANT). Currently, the function of this protein and its homologs are totally unknown. Meanwhile, the homologs of ORF3141 are mainly present in high GC Gram-positive bacteria. We proposed that the secreted protein of ORF3141 was possibly a virulence factor of *N*. *seriolae* strain ZJ0503. In this study, the function of ORF3141 was explored and whether this protein was the bacterial virulence factor that induced infection in fish was determined. Gene cloning, secreted protein identification, subcellular localization, overexpression, and apoptosis detection assays of ORF3141 and ORF3141Δsig proteins were performed. Moreover, the results indicated that *ORF3141* gene encoded a secreted protein which contained a mitochondrial targeting sequence was and involved in apoptosis regulation. These results may lay the foundation for further study on the function of ORF3141 protein and promote the understanding of the molecular pathogenic mechanism of *N*. *seriolae*.

## Results

### Cloning and the sequence analysis of ORF3141

The *ORF3141* and *ORF3141Δsig* genes of *N*. *seriolae* strain ZJ0503 were cloned, and the recombinant plasmids of pEGFP-3141, pEGFP-3141Δsig, pcDNA-3141, and pcDNA-3141Δsig were constructed successfully. Sequence analysis revealed that an open reading frame (ORF) of *ORF3141* gene was 558 bp, and the deduced amino acid sequence had a length of 185 residues (Fig. 1A). The predicted molecular weight of ORF3141 protein was 18.96 kDa, and the theoretical isoelectric point (pI) was 5.85. The instability index of ORF3141 was computed to be 25.43, indicating its stability. The grand average of hydropathicity was -0.112, which signifies that the ORF3141 protein was hydrophilic. ORF3141 was predicted to be a secreted protein with a cleavable signal peptide at N-terminus and co-localized with the mitochondria in the host. ORF3141 was composed of a signal peptide (residues 1-30, MKLLNPRGFGLVCASAAVAAGLMLAGCANT) at N-terminate, two low complexity domains (residues 47-70 and 90-117), and five kinds of functional sites including N-myristoylation, Protein kinase C phosphorylation, Casein kinase II phosphorylation, N-glycosylation, and cAMP- and cGMP-dependent protein kinase phosphorylation sites (Fig. 1B). No protein superfamily members of ORF3141 protein were found. The three-dimensional structure of ORF3141 from *N*. *seriolae* was predicted by I-TASSER. The structure was closed to that of multidrug ABC transporter Sav1866 from *Staphylococcus aureus* (Fig. 1C), which was classified as a hydrolase [27]. Moreover, the protein contained two ligand binding sites (residues 108 and 111) of coenzyme F430 and an enzyme active site (residues 107) of ATP phosphohydrolase (steroid-exporting).

**Fig 1.**
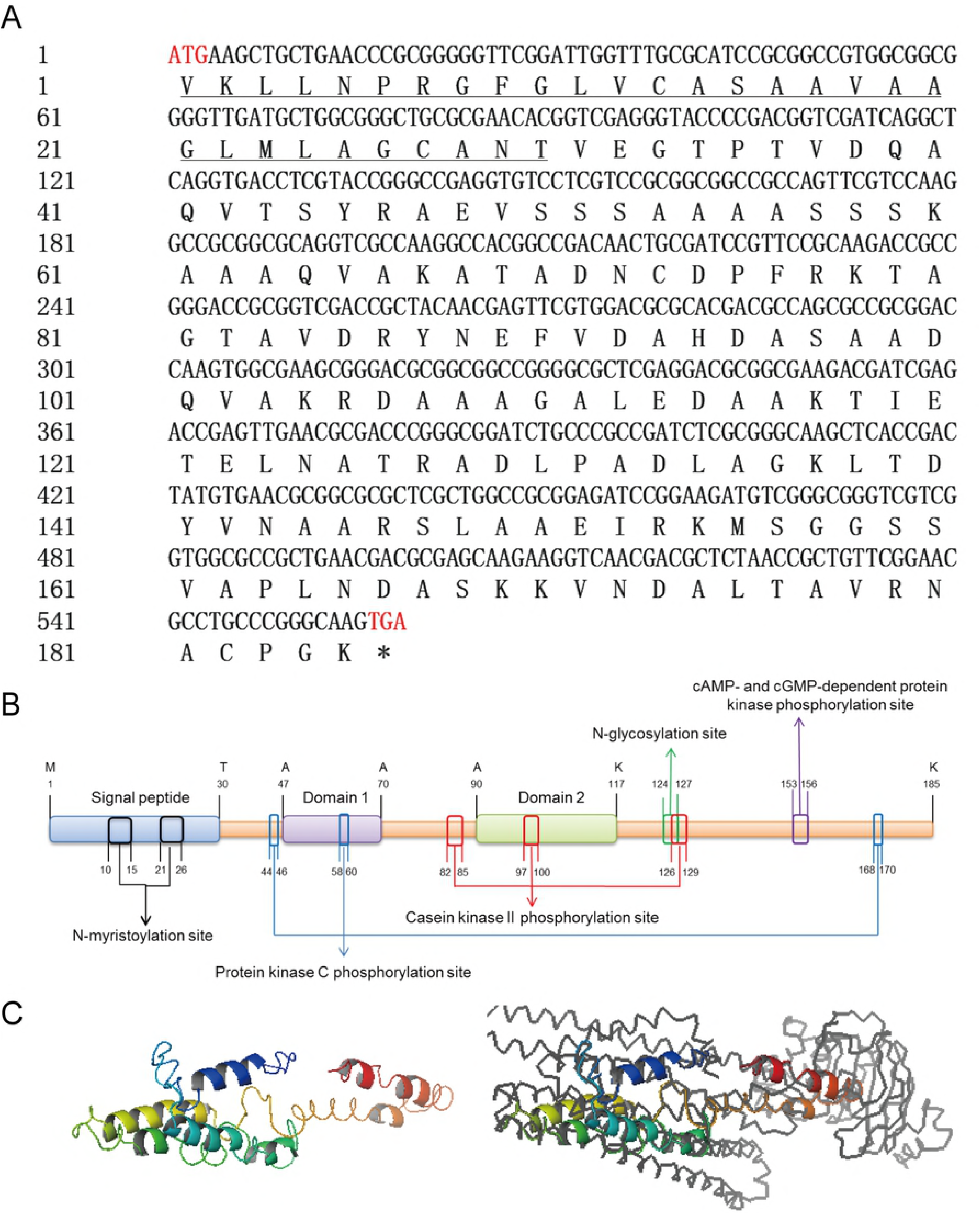
Sequence and structure analysis of ORF3141 from *N*. *seriolae*. (A) The full lengths nucleotide and deduced amino acid sequence of *ORF3141* gene. The underline amino acid sequence showed the signal peptide. (B) Schematic representation of the domain topology of ORF3141. The ORF3141 protein was comprised of a signal peptide at N-terminate (residues 1-30), two low complexity domains (residues 47-70, 90-117), and five kinds of functional sites. (C) The three-dimensional structures of ORF3141 protein from *N*. *seriolae* (Left) and the closest structure to multidrug ABC transporter Sav1866 from *Staphylococcus aureus* (Right). The diagrams were generated using I-TASSER online and PyMOL 1.8 software.

Protein BLAST showed that the deduced amino acid sequence of ORF3141 protein displayed low identity with homologous sequence from Actinomycetes, with the highest identity of *N. concava* (84%) and the lowest identity of *Mycobacterium marinum* (23%). This protein had a rather high homology in *Nocardia* species (61% - 84%) (S1 Table) and its sequence was also found in other *N*. *seriolae* stains, such as *N*. *seriolae* N-2927 [28], *N*. *seriolae* U-1 [29], *N*. *seriolae* EM150506, *N*. *seriolae* CK-14008 and *N*. *seriolae* UTF-1 [30] (S2 Table). Multiple alignment of the deduced amino acid sequences of the ORF3141 protein showed no putative conserved domains (Fig. 2A). A phylogenetic tree was constructed to identify the evolutionary relationships between *N*. *seriolae* with other species of ORF3141 homologs, on the basis of the deduced amino acid sequence of ORF3141. The results showed that ORF341 proteins in *Nocardia sp*. were clustered in one group (Fig. 2B).

**Fig 2.**
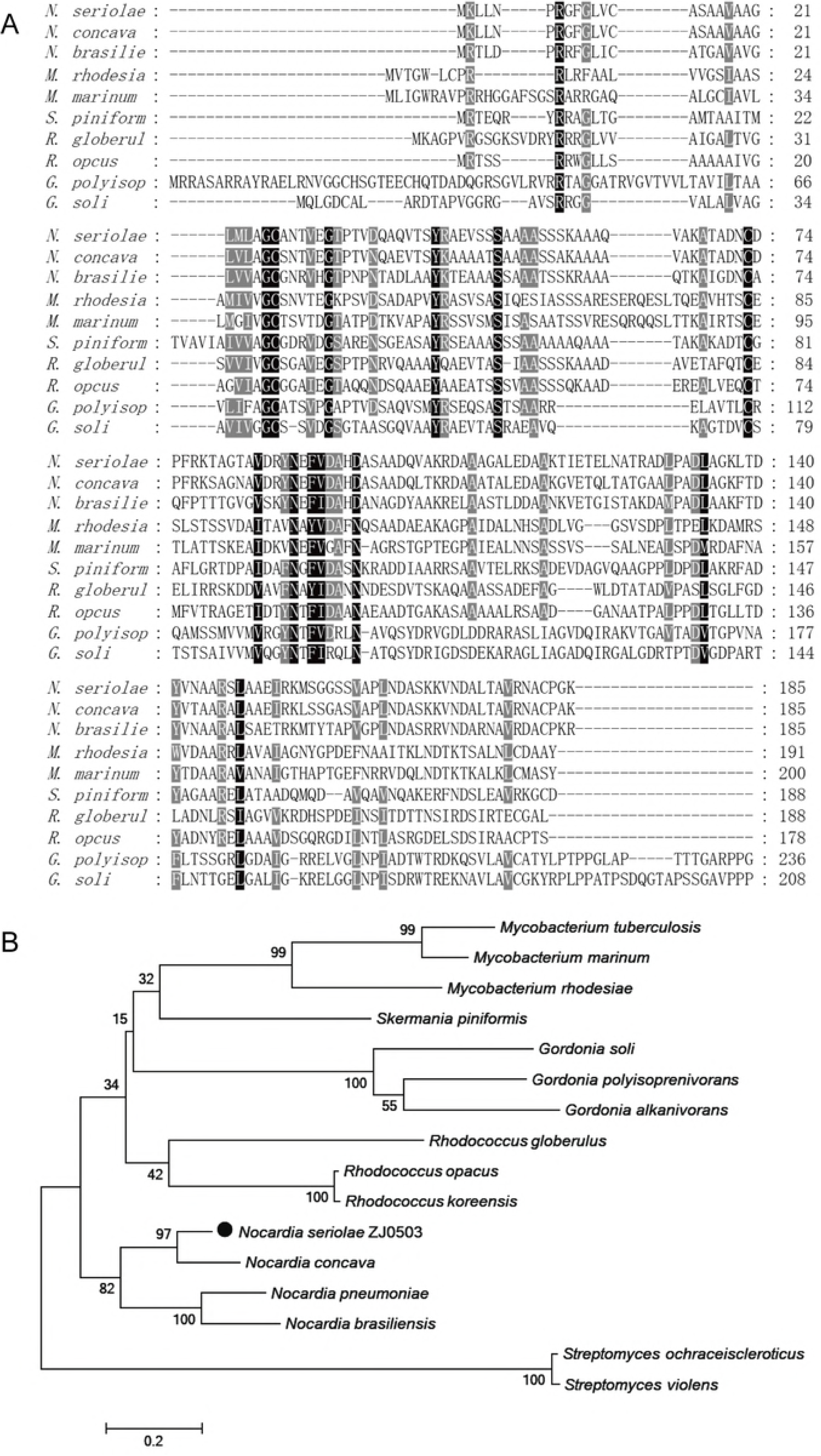
Multiple sequence alignment and construction of phylogenetic tree. (A) Multiple alignment of the deduced amino acid sequences of ORF3141 protein among different species. Shaded regions indicate residues sharing homology, black regions indicate 100 % homology, dark gray regions indicate homology higher than 75 %. (B) Construction of phylogenetic tree among *N*. *seriolae* and other species with ORF3141 protein homology sequences. Protein sequences were aligned with Clustal W, and the nonrooted neighbor-joining tree was generated by MEGA 5.0 program. Number at branch points indicate bootstrap support. GenBank accession numbers are shown as followed: *Mycobacterium tuberculosis* H37Ra (ABQ74022.1), *Mycobacterium marinum* M (ACC41759.1), *Mycobacterium rhodesiae* NBB3 (AEV74934.1), *Skermania piniformis* (WP_066468955.1), *Gordonia soli NBRC* 108243 (GAC69437.1), *Gordonia polyisoprenivorans* NBRC 16320 (GAB24816.1), *Gordonia alkanivorans* NBRC 16433 (GAA10605.1), *Rhodococcus globerulus* (WP_045067034.1), *Rhodococcus opacus* (WP_012688156.1), *Rhodococcus koreensis* (WP_072945555.1), *Nocardia seriolae* ZJ0503 (WP_033086993.1), *Nocardia concave* (WP_040804030.1), *Nocardia pneumonia* (WP_040776180.1), *Nocardia brasiliensis* NBRC 14402 (GAJ80338.1), *Streptomyces ochraceiscleroticus* (WP_031055338.1), *Streptomyces violens* (WP_030248901.1).

### Identification of ORF3141 protein was a secreted protein

Preparation and shotgun mass spectrometry (MS) of the extracellular products of *N*. *seriolae* stain ZJ0503 showed that the characteristic peptide “TIETELNATR” of ORF3141 was detected with confidence greater than or equal to 99%. This result confirmed that ORF3141 was a secreted protein of *N*. *seriolae*.

### Subcellular localization of ORF3141 and ORF3141Δsig proteins in FHM cells

Subcellular localization of ORF3141 and ORF3141Δsig proteins in the FHM cells were determined by the expression of ORF3141-GFP and ORF3141Δsig-GFP fusion protein, respectively. ORF3141-GFP and ORF3141Δsig-GFP fusion protein were detected with strong green fluorescence signal at 48 h post-transfection (hpt). The mitochondria were shown with red fluorescence, and the nucleus were displayed with blue fluorescence. Compared with the location of mitochondria and nucleus in the pEGFP-3141 or pEGFP-3141Δsig transfected cells, the ORF3141-GFP fusion protein was exhibited an aggregated distribution and co-localized with the mitochondria in the cytoplasm, while the ORF3141Δsig-GFP fusion protein was distributed in the whole FHM cell and did not co-localize with the mitochondria. These contrasting results indicated that the N-terminate signal peptide played an important role in mitochondrial targeting. In addition, the expression of ORF3141 or ORF3141Δsig proteins changed the distribution of the mitochondria from perinuclear halo into lumps, and the apoptotic bodies were detected. By contrast, the signal of GFP was distributed in both cytoplasm and nucleus in the control cells which were transfected with pEGFP-N1, and no specific fluorescence co-localized with the mitochondria (Fig. 3).

**Fig 3.**
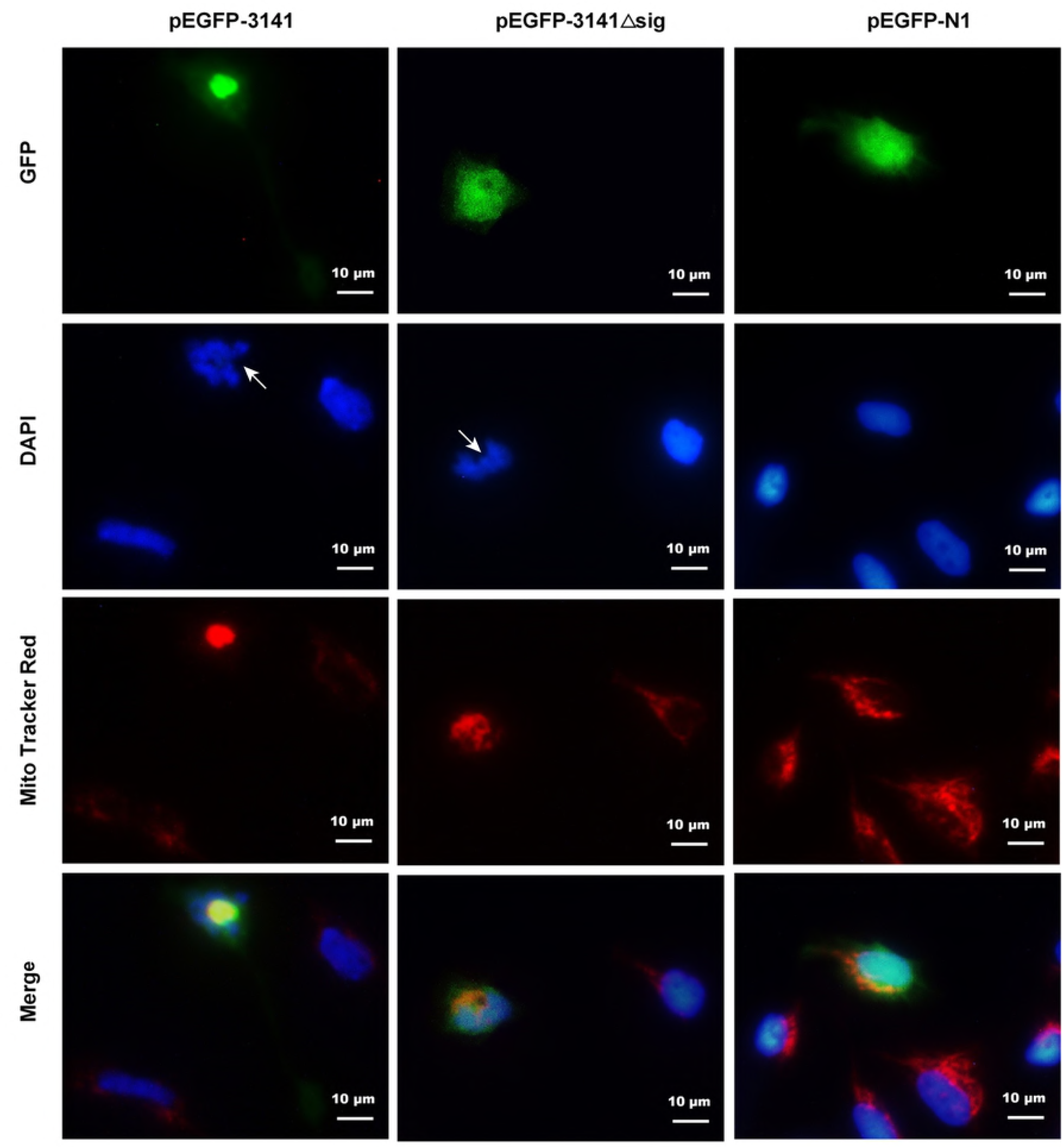
Subcellular localization of ORF3141 and ORF3141Δsig proteins in FHM cell. Green fluorescence shows the ORF3141-GFP, ORF3141Δsig-GFP or GFP, red fluorescence shows the mitochondria, blue fluorescence shows the nucleus, and the arrow refers to the nuclear fragmentation and condensation. In the control plasmid pEGFP-N1, the green fluorescence was evenly distributed in the whole cell of FHM cells, mitochondria were annularly distributed in the periphery of nucleus, the edge of nucleus was smooth with uniform dyeing and had no apoptosis characteristics. Subcellular localization of ORF3141-GFP fusion proteins were mainly distributed in the cytoplasm that coincided with the distribution of mitochondria, which indicated that the protein ORF3141 is targeted to mitochondria. Differently, subcellular localization of ORF3141Δsig-GFP fusion proteins were evenly distributed in the whole cell of FHM cells, and were not coincide with the distribution of mitochondria, which indicated that the subcellular localization changed with the excision of signal peptide. The expression of ORF3141 or ORF3141Δsig proteins changed the distribution of mitochondria from perinuclear halo into lumps and apoptotic bodies were detected.

### Apoptosis induced in FHM cells by overexpression of ORF3141 and ORF3141Δsig proteins

To investigate whether ORF3141 and ORF3141Δsig proteins were involved in the apoptosis of fish cells, plasmids pcDNA-3141 and pcDNA-3141Δsig were transfected into the FHM cells, respectively. Several typical apoptotic features, such as cellular shrinkage, blebbing of the nuclear membrane, nuclear pyrosis, and nuclear fragmentation, in ORF3141 and ORF3141Δsig proteins overexpressed FHM cells were observed by microscope with fluorescent and white light (Fig. 4). Thus, the ORF3141 and ORF3141Δsig proteins might lead to fish cell apoptosis. The number of apoptotic bodies were counted and used to calculate the apoptotic rate (number of apoptotic bodies / total number of cells × 100%) [31]. The results showed that 19.82 % and 17.05% cells underwent apoptosis in pcDNA-3141 and pcDNA-3141Δsig transfected groups respectively, while only 2.75% cells in control group (Fig. 5A). At 48 hpt, the expression of ORF3141 or ORF3141Δsig proteins in the plasmids transfected FHM cells were confirmed by the presence of a specific band using RTPCR (Fig. 5B) and Western blot analysis (Fig. 5C).

**Fig 4.**
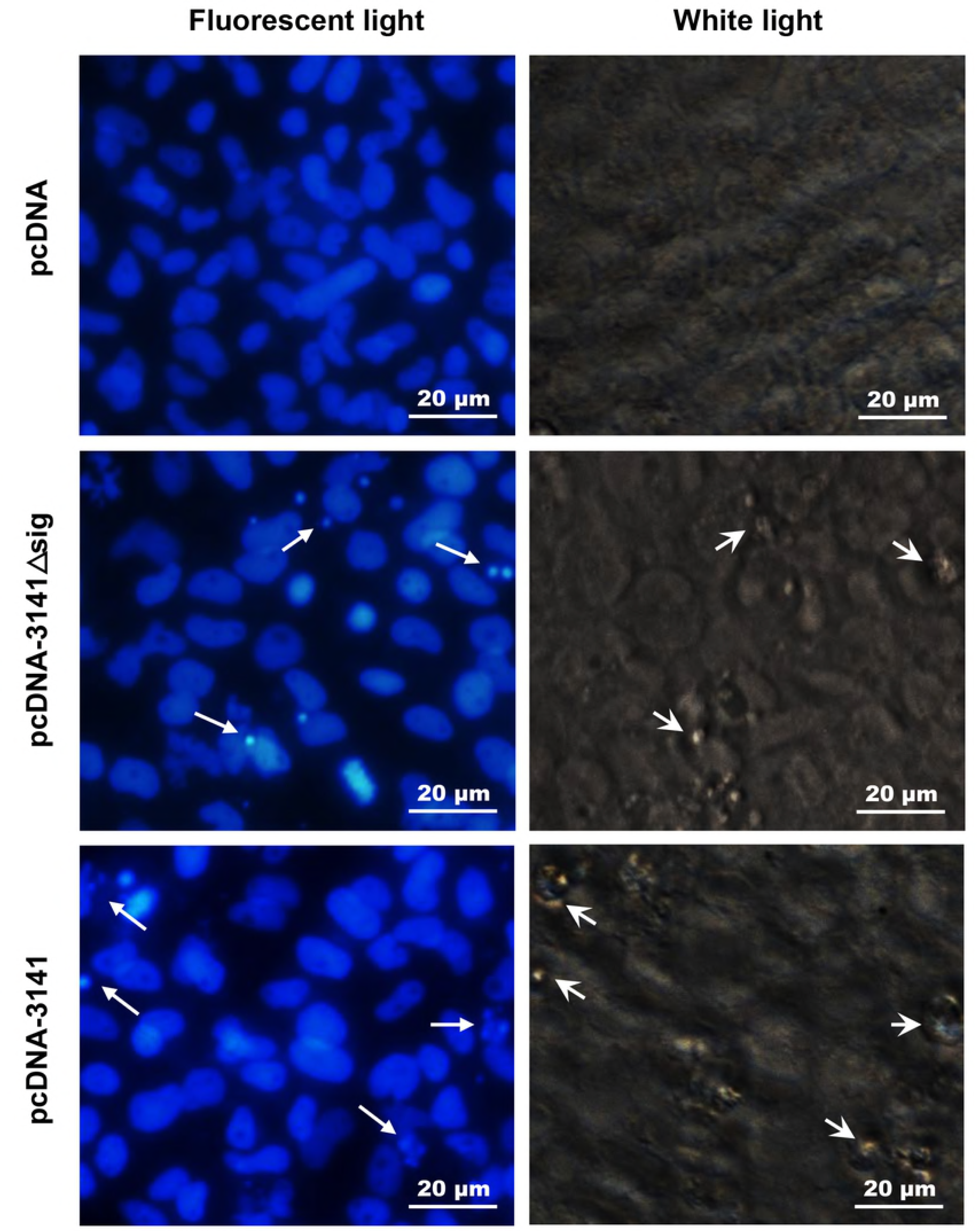
Overexpression of ORF3141 and ORF3141Δsig proteins in FHM cells. The transfected cells were fixed at 48 h post-transfection staining by DAPI. Arrows indicated the apoptotic bodies (fragmented nucleus), arrow heads indicated the apoptotic cells.

**Fig 5.**
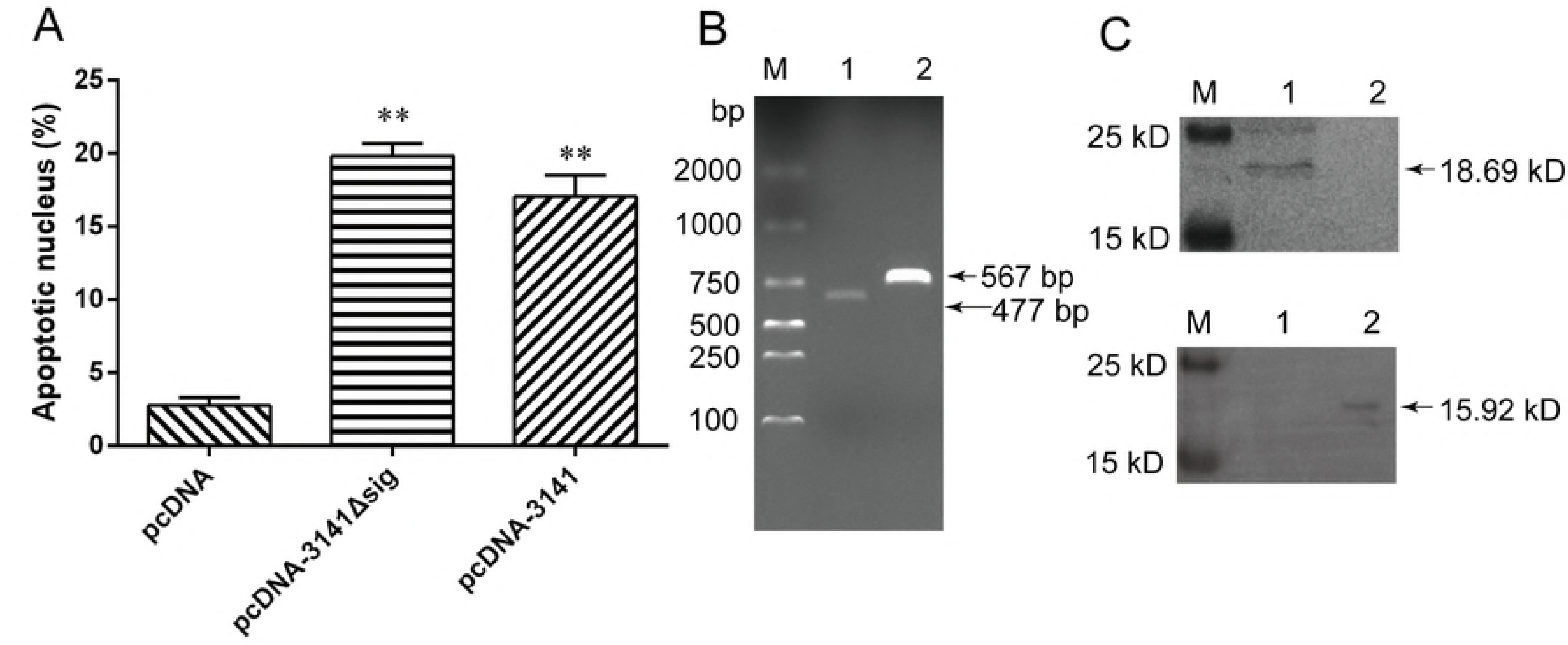
Apoptosis nucleus ratio, RT-PCT, and Western blot of pcDNA-3141 and pcDNA-3141Δsig transfected FHM cells. (A) The apoptotic nucleus was counted and the percentages of apoptotic cells were calculated. Data represent the means for three independent experiments and error bars indicate SD (**p < 0.01). (B) Confirmation of the ORF3141 and ORF3141Δsig expression in FHM cells by RT-PCR. Lane M, DNA marker; lane 1, pcDNA-3141Δsig/FHM; lane 2, pcDNA-3141/FHM. (C) Confirmation of the ORF3141 expression in FHM cells by western blot. Lane M, Protein marker; lane 1, pcDNA-3141/FHM; lane 2, pcDNA 3.1 His/A (Up) and confirmation of the ORF3141Δsig expression in FHM cells by western blot. Lane M, protein marker; lane 1, pcDNA 3.1 His/A; lane 2, pcDNA-3141Δsig/FHM (Down).

### Apoptotic detection of ORF3141 and ORF3141Δsig overexpressed FHM cells

TheΔΨm was shown with the evident damage in JC-1 polymer/monomer fluorescence ratio in ORF3141 or ORF3141Δsig overexpressed cells. TheΔΨm values of both pcDNA-3141 and pcDNA-3141Δsig transfected groups cells were decreased to minimum values at 72 hpt, which were 0.54- and 0.53-folds lower than that of the control cell group, respectively (Fig. 6A). Meanwhile, measurement of caspase-3 activity showed that caspase-3 was activated in ORF3141 or ORF3141Δsig proteins overexpressed cells and reached maximum value at 48 hpt, which were approximately 1.23- and 1.80-folds higher than that of the control cell group, respectively (Fig. 6B). Several apoptosis-related genes of Bcl-2 family mRNA expression were investigated at 0, 24, 48, 72, and 96 hpt by qRT-PCR, including B cell lymphoma-2 (*Bcl-2*), Bcl-2 associated X (*Bax*), Bcl-2 antagonist of cell death (*Bad*), and Bcl-2 interacting domain death agonist (*Bid*) genes. The results showed that pro-apoptotic (*Bad*, *Bid*, and *Bax*) and anti-apoptotic (*Bcl-2*) genes were rapidly activated in both pcDNA-3141 and pcDNA-3141Δsig transfected groups cells and theirs mRNA expression reached peak level at 72 hpt, respectively. Moreover, there were 11.19- and 10.60-folds of *Bax* to *Bcl-2* genes mRNA expression in pcDNA-3141 and pcDNA-3141Δsig transfected groups higher than that of 5.17-fold in the control group at 72 hpt (Fig. 6C). All quantitative assays ofΔΨm value, caspase-3 activity, and apoptosis-related genes mRNA expression indicated that cell apoptosis could be induced by the overexpression of ORF3141 or ORF3141Δsig proteins in the FHM cells.

**Fig 6.**
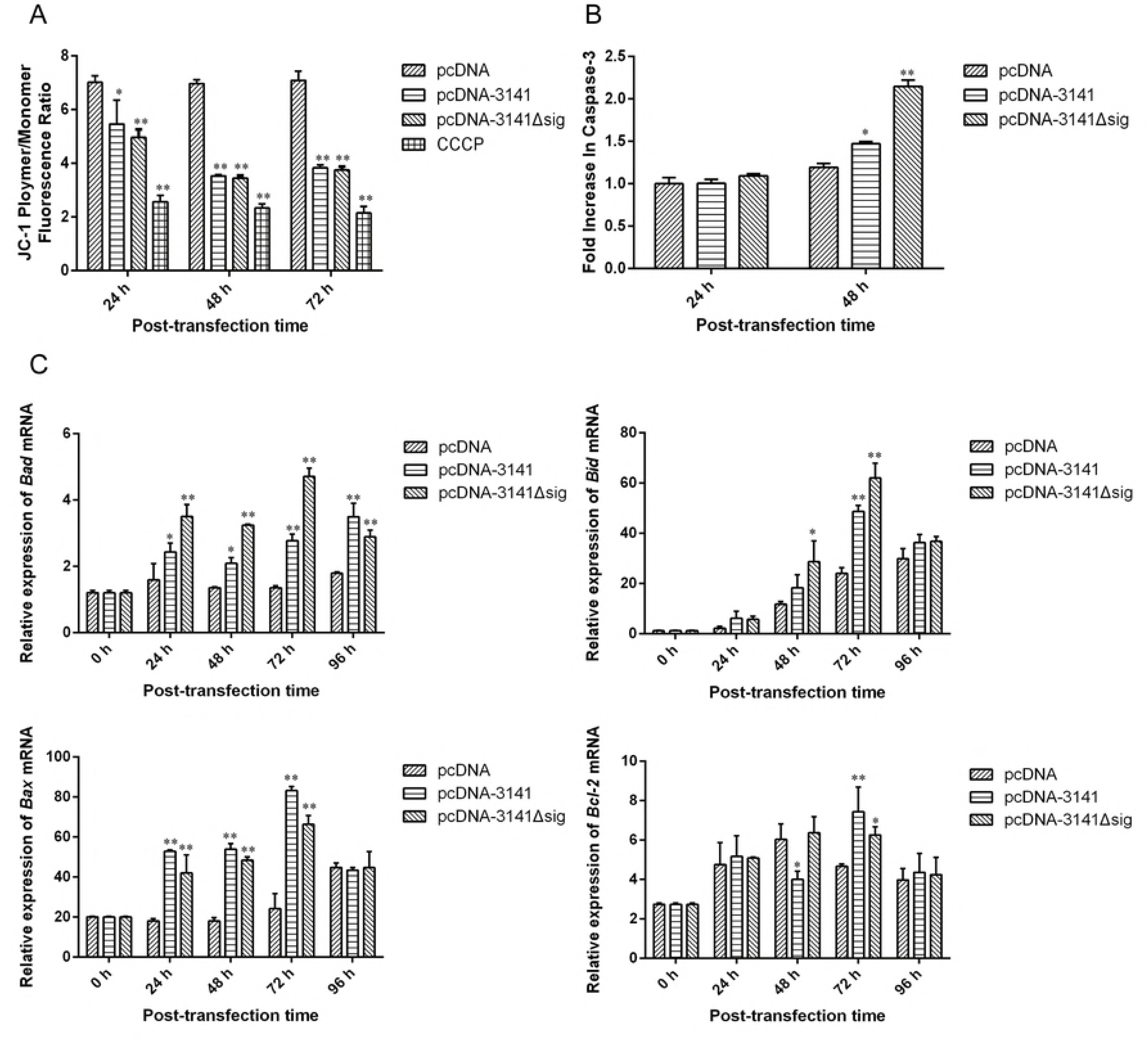
Apoptotic detection of ORF3141 and ORF3141Δsig overexpressed FHM cells. (A) Mitochondrial membrane potential assay. FHM cells transfected with pcDNA-3141, pcDNA-3141Δsig or pcDNA 3.1 His/A plasmid were collected at indicated time points after transfection and the mitochondrial membrane potential were assessed using the JC-1. Un-transfected cells treated with CCCP was positive control. The data were expressed as the JC-1 polymer/monomer fluorescence ratio and error bars indicate SD (\**p* < 0.05, \*\**p* < 0.01). (B) Measurement of caspase-3 activity. FHM cells transfected with pcDNA-3141, pcDNA-3141Δsig or pcDNA 3.1 His/A plasmid were collected at indicated time points after transfection and the levels of cleaved caspase-3 were measured. The data were expressed as fold increase compared to the corresponding caspase-3 activity values in un-transfected cells and error bars indicate SD (\**p* < 0.05, \*\**p* < 0.01). (C) qRT-PCR analysis of the expression of apoptosis-related genes (*Bad*, *Bid*, *Bax*, and *Bcl-2*) and error bars indicate SD (\**p* < 0.05, \*\**p* < 0.01).

## Discussion

No relevant literature on the ORF3141 protein of *Nocardia* or other species is available, and this protein remains largely unknown. Here, we preliminary investigated the characteristic, structure, and function of ORF3141 from *N*. *seriolae* strain ZJ0503. Moreover, *ORF3141* and *ORF3141Δsig* were successfully cloned. The sequence analysis showed an ORF of *ORF3141* gene of 558 bp. This gene encoded a protein with 185 amino acid residues, molecular weight of 18.96 kDa, and isoelectric point of 5.85. A phylogenetic tree based on the deduced amino acid sequence of ORF3141 shared low homology between *N*. *seriolae* and other homologous sequences from Actinomycetes. However, ORF3141 had a rather high homology in *Nocardia* species. Bioinformatic analysis revealed that ORF3141 had no putative conserved domains and no superfamily. This protein comprised a signal peptide and two low complexity domains, but the function of these domains remains unknown. *ORF3141* was predicted to encode a secreted protein and co-localized with the mitochondria in the host cells. Experimentally, the MS analysis of the extracellular products from *N*. *seriolae* strain ZJ0503 proved that ORF3141 was a secreted protein. Subcellular localization of ORF3141 in the host cells exhibited an aggregated distribution and coincided with the mitochondria. Thus, ORF3141 co-localized with the mitochondria in the host cells. Distinguishingly, ORF3141Δsig protein was evenly distributed in the whole cell and did not co-localize with the mitochondria in the host cells. These contrasting results were due to the presence or absence of the signal peptide. Similar results were found in the subcellular localization study of EGFP-GRP75 fusion proteins; that is, GRP-75 with signal peptide was exclusively co-localized with the mitochondria, whereas GRP-75 without the signal peptide was distributed in the whole host cell [32]. Thus, the N-terminate signal peptide was important for mitochondrial targeting. The mitochondrial targeting protein contains specific sequences with specific information. Five kinds of mitochondrial targeting signals (MTSs) are used to determine the protein localization with the mitochondrial outer membranes, inner membranes, or matrix [33]. Many proteins are destined for the mitochondrial matrix typically cleavable targeting signal at the N-terminate [34], which are commonly 20 - 60 amino acids in length and are cleaved upon import into the mitochondrial matrix by the mitochondrial processing peptidase [35,36]. Therefore, ORF3141 was assumed to target the mitochondria matrix.

The function of ORF3141 has not been studied, until now. In our study, we observed that the distribution of mitochondria was altered from perinuclear halo into lumps, and apoptotic bodies appeared during the subcellular localization and overexpression assays, which indicated that cell apoptosis was induced after ORF3141 or ORF3141Δsig proteins stimulating in FHM cells. Mitochondria appear to play a central role in the regulation of several cell death pathways, such as apoptosis, autophagic cell death, necrosis, pyroptosis, and pyronecrosis [37,38]. Other characterized bacterial proteins had also been shown to target the mitochondria and induce apoptosis. For example, Omp38 of *Acinetobacter baumannii* was shown to target mitochondria of Hep-2 cells where it induced the release of Cyt c and AIF, to promote apoptosis [39]. When expressed in human cells, PorB of *Neisseria gonorrhoeae* co-localized with the mitochondria and caused dissipation ofΔΨm but did not induce the release of Cyt c [40]. SipB of *Salmonella enterica* targeted the host cell mitochondria during infection and supported autophagy-mediated cell death in caspase-1 deficient macrophages [41]. As a mitochondrial targeting bacterial secreted protein, ORF3141 might participate in the regulation of cell apoptosis. Distinctively, ORF3141Δsig protein was observed to also induce cell apoptosis without targeting the mitochondria in FHM cells. Thus, it was proved that the domain part without the signal peptide of ORF3141 protein had a critical relationship with cell apoptosis whether it was targeted to the mitochondria or existent in the cytoplasm. Moreover, quantitative assays ofΔΨm value, caspase-3 activity, and apoptosis-related genes mRNA expression were performed to determined how did the ORF3141 and ORF3141Δsig protein participate in cell apoptosis in FHM cells. The Bcl-2 family proteins are critical regulators of mitochondrial apoptosis, which include the anti-apoptotic multidomain members (containing all four BH domains), the pro-apoptotic multidomain members (containing three BH domains), and pro-apoptotic BH3-only members (containing the sole BH3 domain). In addition, a large number of Bcl-2 family protein contain a hydrophobic transmembrane anchoring (TM) domain at the C-terminus which allows them to co-localize to subcellular membranes [42,43]. Among these proteins, Bcl-2 was discovered as an anti-apoptosis protein that blocks cell apoptosis, while Bax was clarified as a pro-apoptosis protein which can form large openings in lipid bilayers to control the MOMP and induce apoptosis [44–46]. Thus, a slight change in the dynamic balance of Bax and Bcl-2 seem to determine the inhibition or promotion of cell apoptosis with apoptotic stimuli [47]. In fact, it is clear that Bcl-2 family proteins can also localize to cytosol and other organelles, including the Golgi apparatus, the endoplasmic reticulum (ER), and the nuclear out membrane (NOM) or the nucleus itself [48]. In non-apoptotic cells, Bax localization is essentially diffuse in the cytosol but it is relocated to mitochondria after cell apoptosis induction [49]. Therefore, it could be speculated that both ORF3141 and ORF3141Δsig proteins were initiated by activating the apoptosis-related genes (Bad, Bid, Bax, and Bcl-2) and induced the translocation of Bax from the cytosol to the mitochondria in FHM cells. Besides, the relative ratio of *Bax* and *Bcl-2* genes mRNA expression was out-of-balance, which were responsible for theΔΨm values declined and caspase-3 activity increased in this study. Although the subcellular localization of ORF3141 and ORF3141Δsig proteins were different, they could induce the cell apoptosis via the mitochondrial pathway.

Further studies are required to verify the mechanisms involved in ORF3141-induced cell apoptosis by yeast two hybrid experiment. Whether the ORF3141 was the major virulence factor of *N*. *seriolae* remains to be clarified by constructingΔORF3141 mutant attenuated and ORF3141 mutant completed *N*. *seriolae*. The relationship of the interaction between ORF3141 and macrophage also need to be highlighted in future studies. In summary, we cloned an *ORF3141* gene from *N*. *seriolae* strain ZJ0503 and analyzed its sequence structure. Moreover, MS analysis, subcellular localization, overexpression assay, and apoptosis detection were performed. These results showed that ORF3141 of *N*. *seriolae* strain ZJ0503 was a secreted protein that targeted the host cell mitochondria and induced apoptosis in transfected FHM cells.

## Materials and methods

### Bacterial strains, FHM cells, and plasmids

*N*. *seriolae* strain ZJ0503 was isolated from diseased golden pompano (*T*. *ovatus*) in YangJiang City, Guangdong Province, China [26] and was cultured in an optimized medium [glucose 20 g·L^-1^, yeast extract 15 g·L^-1^, K_2_HPO_4_ 0.75 g·L^-1^, CaCl_2_ 0.2 g·L^-1^ (sterilized separately), and NaCl 5 g·L^-1^, PH 6.5 ± 0.2)] at 28 °C [50]. *Escherichia coli* DH5α was used for gene cloning and it grown in Luria-Bertani (LB) medium with vigorous shaking at 37 °C. Fathead minnow (FHM) epithelial cells [51] were stored at -196 °C in Guangdong Provincial Key Laboratory of Pathogenic Biology and Epidemiology for Aquatic Economic Animals and cultured at 25 °C in Leibovitz’s L15 medium containing 10% fetal bovine serum (Invitrogen, USA) in 5% CO_2_ incubator. Plasmid pEGFP-N1 was used for subcellular localization, while pcDNA3.1/His A was used for overexpression.

### Ethics statement

All animal experiments were handled in accordance with guidelines defined by the Institutional Animal Care Use Committee (IACUC) of Guangdong Ocean University and were approved by Guangdong Provincial Key Laboratory of Pathogenic Biology and Epidemiology for Aquatic Economic Animals, following strict compliance with the regulations of the local government.

### Gene cloning and plasmid construction of *ORF3141* and *ORF3141Δsig*

Genomic DNA was extracted from *N*. *seriolae* strain ZJ0503 using TIANamp Bacteria DNA Kit (Tiangen, Beijing) following the manufacture’s instruction. Four pairs of different primers were carefully designed with corresponding restriction enzyme sites using Primer 5.0, on the basis of *ORF3141* and *ORF3141*Δsig genes of the whole genome sequence data of *N*. *seriolae* strain ZJ0503. The PCR primers of pEGFP-3141F/R and pcDNA-3141F/R (Table 3) were used to amplify the *ORF3141* gene. The PCR primers of pEGFP-3141ΔsigF/R and pcDNA-3141ΔsigF/R (Table 1) were used to amplify the *ORF3141Δsig* gene. The PCR procedures were performed with KOD-plus-Neo DNA polymerase (Toyobo, Osaka, Japan), using the following PCR program: pre-denaturation at 98 °C for 2 min, 30 cycles at 98 °C for 10 s, 55 °C for 15 s, 69.5 °C for 15 s; and a final extension at 68 °C for 5 min. All PCR products of *ORF3141* and *ORF3141Δsig* genes were electrophoresed on 1% agarose gel and purified using EasyPure PCR Purification Kit (TRANSGEN, Beijing). The purified PCR product of *ORF3141* was digested by corresponding restriction enzymes, ligated into eukaryotic vectors pEGFP-N1 and pcDNA3.1/His A, respectively. And then transformed into competent *E*. *coli* DH5α cells. The different constructs were confirmed by corresponding restriction enzyme digestion and DNA sequencing by Guangzhou Sangon Biologic Engineering & Technology and Service Co. Ltd. The purified PCR product of *ORF3141Δsig* was processed similarly. Finally, the constructed recombinant plasmids were named as pEGFP-3141, pEGFP-3141Δsig, pcDNA-3141, and pcDNA-3141Δsig.

**Table 1.**
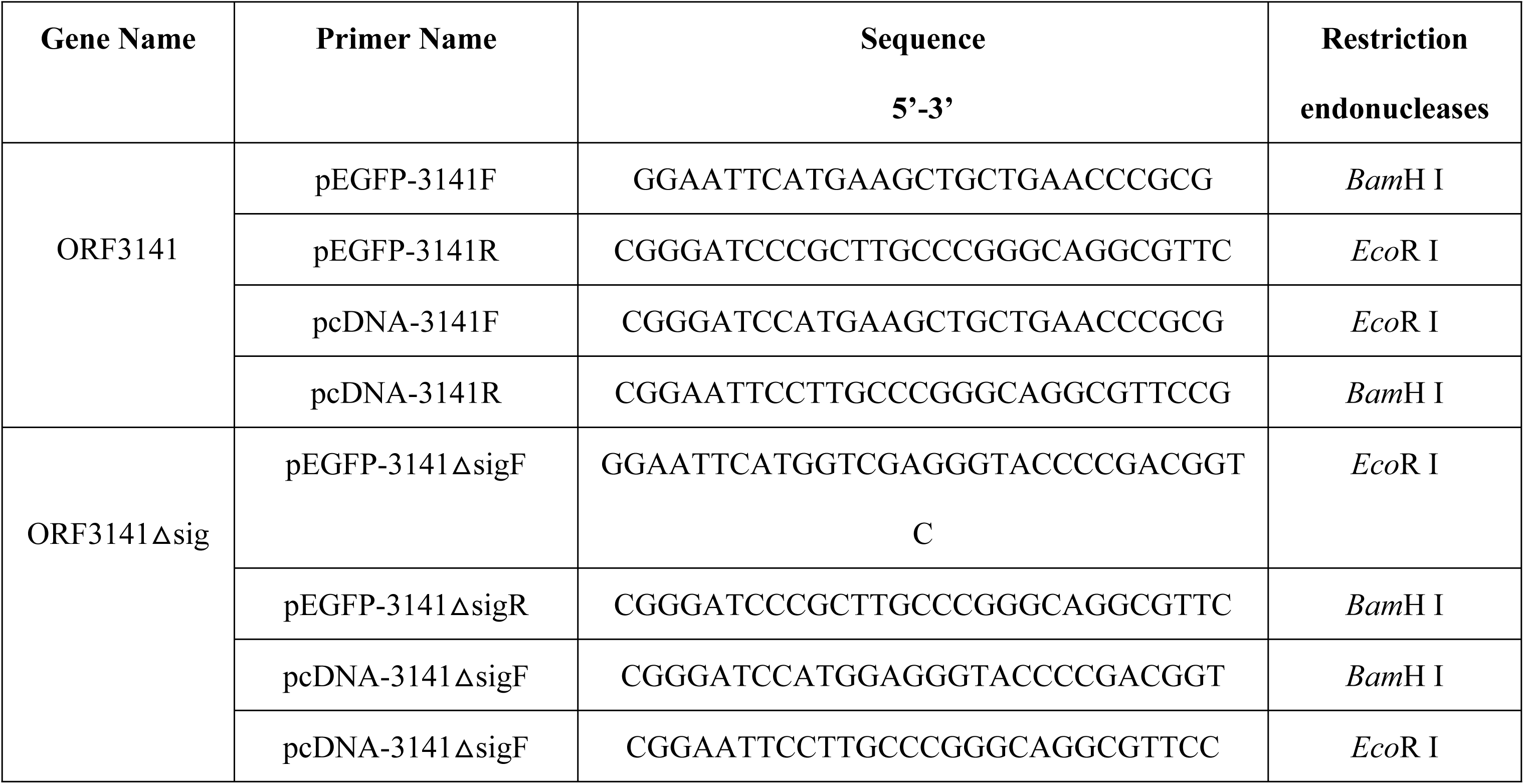
Primers used for gene cloning

### Bioinformatics analysis, sequence alignments, and phylogenetic analysis

Based on the whole genome sequence data of *N*. *seriolae* strain ZJ0503, sequence analysis was performed with the BLAST program using NCBI (http://www.ncbi.nlm.nih.gov/BLAST/). The amino acid sequence for ORF3141 was deduced, and the physical and chemical properties were predicted using ExPASy software (http://www.expasy.org/). The location of domains was predicted by the InterProScan program (http://www.ebi.ac.uk/Tools/pfa/iprscan/). The typical structures of ORF3141 protein were predicted by I-TASSER online software (http://zhanglab.ccmb.med.umich.edu/I-TASSER/). The potentially excreted proteins were predicted by LocTree3 (https://rostlab.org/services/loctree/) and ExPASy-PROSITE. The subcellular localization and signal peptides were predicted with LocTree 3 and SignalP 4.1 Server (www.cbs.dtu.dk/services/SignalP/), respectively. The MTSs were predicted using TarfetP (http://www.cbs.dtu.dk/services/TargetP/) and PSORT II (https://psort.hgc.jp/form2.html). Protein multiple sequence alignments of ORF3141 protein were performed by ClustalX 2.0 program with the default parameters and edited by the GeneDoc software. The phylogenetic tree was generated based on the deduced amino acid sequence of ORF3141 with the neighbor-joining method using MEGA 6 program, in which the Poisson distribution substitution model and bootstrapping procedure with 1000 bootstraps were applied.

### Preparation and identification of the *N. seriolae* extracellular products

The extracellular products of *N*. *seriolae* strain ZJ0503 were obtained by cellophane overlay method [52]. Specifically, *N*. *seriolae* were grown on optimized medium agar plate at 28 °C for 2 d, and a single colony was prepared for bacterial suspension. Then, 100 μL of the bacterial suspension was spread closely on optimized medium plates covered with sterile cellophane sheet and incubated at 28 °C for 3 - 5 d. The *N*. *seriolae* cells grown on the cellophane sheet were stripped away from the optimized medium plates, and the extracellular products were washed down with sterilized PBS subsequently. The harvested suspension was centrifuged at 8000×g, 4 °C for 20 min, and the supernatant containing extracellular products was filter sterilized with a 0.2 μm membrane filter. Then, the sterilized supernatant was transferred into a dialysis tubing (3.5k MW) and dialyzed in ultrapure water at 4 °C for 16 - 24 h. During dialysis, the ultrapure water was changed 3 - 4 times. The purified supernatant was transferred into a centrifuge tube after dialysis and frozen under -80 °C. Finally, it was lyophilized using a vacuum freeze dryer to obtain the protein dry powder which was identified using shotgun MS.

### FHM cell mediated transient expression and subcellular localization

Plasmids of pEGFP-3141, pEGFP-3141Δsig, and pEGFP-N1 were prepared in advance using an endotoxin-free plasmid purification kit (Qiagen Inc., Chatsworth, CA). Subcellular localization of ORF3141 and ORF3 141Δsig proteins were performed through pEGFP-N1 fusion protein expression. Given that a permanent FHM cell line is presently available, FHM cells were cultured in 24-well plates and grown to 70 % confluency for transfection. The FHM cells were transfected with empty pEGFP-N1 plasmids, recombinant pEGFP-3141 plasmids, and recombinant pEGFP-3141Δsig plasmids using Lipofectamine 2000 (Invitrogen, Carlsbad, CA) according to the manufacturer’s protocol, respectively. At 48 hpt, the FHM cells were washed with PBS (pH 7.4), stained with 300 nM MitoTracker Red CMXRos dye (Molecular Probes, Carlsbad, CA) at 28 °Cor 45 min, fixed with 4% paraformaldehyde for 30 min, and labeled with 1 μg/mL diamidino-2- phenylindole (DAPI) at room temperature for 10 min. Finally, the cells were rinsed with PBS and mounted with 50% glycerol. The fluorescence exhibited by the transfected FHM cells were observed using a fluorescence microscope (Leica DM IRB).

### Overexpression of ORF3141 and ORF3141Δsig proteins in FHM cells

The extraction and transfection of recombinant pcDNA-3141 plasmids, recombinant pcDNA-3141Δsig plasmids, and control pcDNA3.1/His A plasmid were performed with reference to step 2.5. The transfected FHM cells were stained with DAPI at 48 hpt and microscopically observed to test whether the overexpression of ORF3141 and ORF3141Δsig proteins induced apoptosis in fish cells. Meanwhile, to identify the expression of ORF3141 and ORF3141Δsig proteins in the transfected FHM cells, the cells were harvested at 48 hpt to extract the total RNA and proteins for RT-PCR and Western blot analysis, respectively. RT-PCR was performed with primers pcDNA-314 1 F/R and pcDNA-3141ΔsigF/R (Table 3) following the synthesis of cDNA. Western blot was conducted using mouse anti-His monoclonal antibody (Sigma, St. Louis, MO) as the primary antibody at a dilution of 1:1000 and horseradish peroxidase-conjugated goat anti-mouse IgG (Sigma, St. Louis, MO) as the secondary antibody at a dilution of 1:5000.

### Quantitative analysis ofΔΨm value, caspase-3 activity and apoptosis-related gene mRNA expression

Several quantitative assays of mitochondrial membrane potential (ΔΨm) value, caspase-3 activity, and apoptosis-related gene mRNA expression (pro-apoptotic gene: *Bad*, *Bid*, and *Bax*; antiapoptotic gene: *Bcl*-*2*), were performed to confirm the apoptosis caused by overexpression of proteins in the transfected recombinant pcDNA-3141 or pcDNA3141Δsig in the FHM cells. TheΔΨm values were assessed at 24, 48, and 72 hpt using the JC-1 assay kit (Beyotime, Shanghai, China) by the method described previously with minor modification [53]. The positive control was treated with carbonyl cyanide 3-chlorophenylhydrazone (CCCP, 10 mM) to medium at 1:1000 25 °C for 20 min. Then,ΔΨm value was measured by changes in the 590/530 JC-1 emitted fluorescence with an Enspire 2300 Multilabel Reader (Perkin Elmer, MA, USA). The caspase-3 activity was detected in the FHM cells at 24 and 48 hpt as described previously [54] using a caspase-3 colorimetric assay kit (BioVision, Milpitas, CA).

Three groups of transfected FHM cells were harvested at 0, 12, 24, 48, 60, and 72 hpt to extract the total RNA, and quantitative real-time PCR (qRT-PCR) was performed following the synthesis of cDNA. The cell apoptotic effect of post-transfection on the expression of apoptosis-related genes mRNA, was investigated using real-time SYBR green PCR Master Mix on Applied Biosystems 7500 Real-Time PCR system (ABI, USA). Each assay was performed in triplicate with *β*-*actin* gene as the internal control. According to the apoptosis gene sequence, five pairs of specific primer (Table 2) were carefully designed for qRT-PCR investigation. The PCR was performed in a 10 μL reaction volume containing 0.5 μL of each primer (10 μM), 0.3 μL of cDNA, 3.7 μL of PCR-grade water, and 5μL of SYBR^®^ Select Master Mix (ABI, USA) according to the manufacturer’s protocol. The PCR conditions were as follows: 95°C for 2 min; 40 cycles of 95 °C for 15 s, and 58 °C for 1 min for four apoptosis-related genes and *β-actin*, respectively. Melt curve analysis of the amplification products was performed over a range of 60 – 95 °C at the end of each PCR reaction to confirm the single product generation. The relative expression levels of four apoptosis-related genes were calculated using the comparative Ct 2^(-ΔΔCt)^ method [55].

**Table 2.**
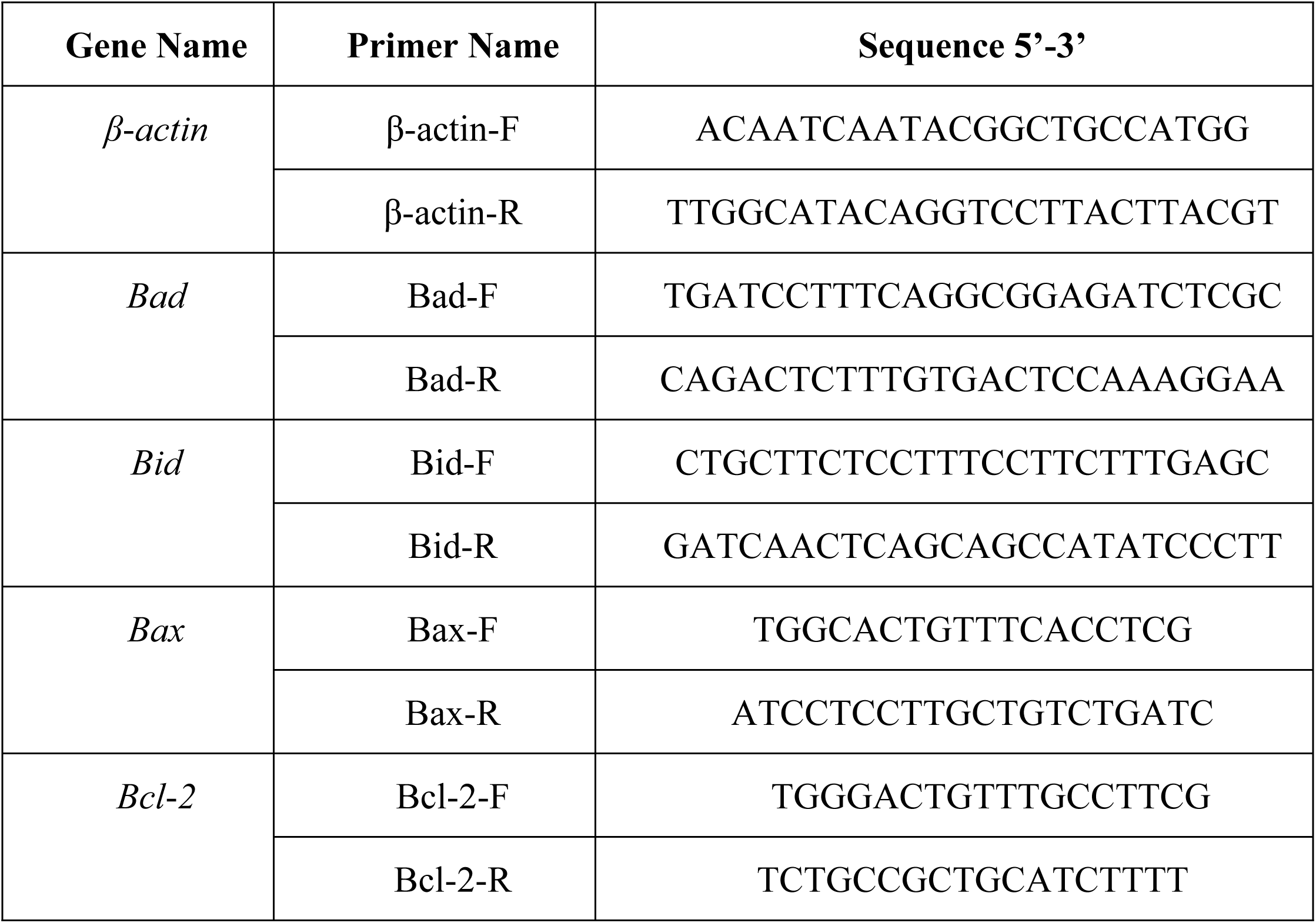
Primes used for apoptosis-related genes investigated by qRT-PCR.

### Statistical analysis

Data were presented as the means ± standard deviation (SD). Statistical analysis was performed with one-way ANOVA with the SPSS statistics 21.0 software and the figures were edited by GraphPad Prism software. Data represent the means for three independent experiments and statistical significant is highlighted with asterisks in the figures as follows: *p* > 0.05, not significant; *p* < 0.05 (*), significant; *p* < 0.01 (**), extremely significant. The means and *p* values for pairwise comparisons of all experiments are provided in S3 Table.

## Acknowledgment

We are grateful to all the laboratory members for the discussion and critical readings of the manuscript.

## Supporting information

**S1 Table.** Compared with the homology of ORF3141 among other Nocardia species.

**S2 Table.** The same sequence of ORF3141 between different N. seriolae strain.

**S3 Table.** Statistical analysis of all experimental data.

